# Investigating brain mechanisms underlying natural reading by co-registering eye tracking with combined EEG and MEG

**DOI:** 10.1101/2021.07.09.451139

**Authors:** Béla Weiss, Felix Dreyer, Elisabeth Fonteneau, Maarten van Casteren, Olaf Hauk

## Abstract

Linking brain and behavior is one of the great challenges in cognitive neuroscience. Ultimately, we want to understand how the brain processes information to guide every-day behavior. However, most neuroscientific studies employ very simplistic experimental paradigms whose ecological validity is doubtful. Reading is a case in point, since most neuroscientific studies to date have used unnatural word-by-word stimulus presentation and have often focused on single word processing. Previous research has therefore actively avoided factors that are important for natural reading, such as rapid self-paced voluntary saccadic eye movements. Recent methodological developments have made it possible to deal with associated problems such as eye movement artefacts and the overlap of brain responses to successive stimuli, using a combination of eye-tracking and neuroimaging. A growing number of electroencephalography (EEG) and functional magnetic resonance imaging (fMRI) are successfully using this methodology. Here, we provide a proof-of-concept that this methodology can be applied to combined EEG and magnetoencephalography (MEG) data. Our participants naturally read 4-word sentences that could end in a plausible or implausible word while eye-tracking, EEG and MEG were being simultaneously recorded. Eye-movement artefacts were removed using independent-component analysis. Fixation-related potentials and fields for sentence-final words were subjected to minimum-norm source estimation. We detected an N400-type brain response in our EEG data starting around 200 ms after fixation of the sentence-final word. The brain sources of this effect, estimated from combined EEG and MEG data, were mostly located in left temporal lobe areas. We discuss the possible use of this method for future neuroscientific research on language and cognition.

## Introduction

There is an increasing demand in cognitive science for experimental paradigms that tap into cognitive processes under real life conditions, for several reasons: 1) to ensure that theories based on lab data are ecologically valid, 2) to turn lab-based experimental paradigms into real-word applications, 3) to develop experimental paradigms that do not require explicit training of participants on unfamiliar tasks, e.g. in order to test clinical populations. Text reading is a case in point. Most behavioral and in particular neuroscientific studies on reading have used word-by-word paradigms, often referred to as (rapid) serial visual presentation paradigms ((R)SVPs). These paradigms have produced a wealth of data, which are the foundation for influential models of visual word recognition (Balota et al., 2007; Carreiras, Armstrong, Perea, & Frost, 2014; Norris, 2009). However, several authors have pointed out that RSVPs lack essential features of natural reading that can significantly affect the word recognition process, the most important ones being parafoveal preview and saccade-related attention shifts (Degno & Liversedge, 2020; Dimigen, Sommer, Hohlfeld, Jacobs, & Kliegl, 2011; Hutzler et al., 2007; Kornrumpf, Niefind, Sommer, & Dimigen, 2016; Rayner, 1998; Weiss, Knakker, & Vidnyanszky, 2016).

Eye movements during reading are sensitive to a range of psycholinguistic and cognitive variables at the single-word and sentence level (Kliegl, Nuthmann, & Engbert, 2006; Rayner, Pollatsek, & Reichle, 2003), which could potentially reveal insights into various aspects of language processing and cognition. Some authors have described saccadic eye movements as “optimal experiments” that reflect the sampling strategy and prior beliefs of the participants (Friston, Adams, Perrinet, & Breakspear, 2012). The simultaneous recording of eye movements and brain responses during reading would therefore provide a wealth of information about brain dynamics during natural language processing and its possible link to behavior. This would be a milestone for basic research and the development of neurocognitive models of language processing, and might link behavioral patterns to neurological deficits in clinical populations.

Normal readers commonly fixate 3-4 words per second. Unfortunately, the temporal resolution of metabolic neuroimaging methods (such as functional magnetic resonance imaging, fMRI) is too low to separate activation from individual words that close in time, let alone to disentangle processing stages within a fixation. fMRI in combination with eye-tracking can produce meaningful results for example for different word categories, similar to conventional event-related designs (Desai, Choi, Lai, & Henderson, 2016; Schuster, Hawelka, Hutzler, Kronbichler, & Richlan, 2016). However, if it is essential to resolve the activation time course within a fixation duration or for neighboring words in a sentence, then methods with higher temporal resolution are required. Over the last decade, a number of studies have used the high temporal resolution of EEG in combination with eye-tracking to study aspects of reading (Baccino & Manunta, 2005; Dimigen et al., 2011; Henderson, Luke, Schmidt, & Richards, 2013; Himmelstoss, Schuster, Hutzler, Moran, & Hawelka, 2019; Hutzler et al., 2007; Metzner, von der Malsburg, Vasishth, & Rosler, 2016; Weiss et al., 2016). Importantly, several studies compared natural reading with RSVP paradigms, and found significant differences between the two (Blythe, Johnson, Liversedge, & Rayner, 2014; Kornrumpf et al., 2016; Metzner et al., 2016).

However, the co-registration of eye-tracking with combined electro- and magnetoencephalography (EEG and MEG) during natural reading has not been reported yet. The combination of EEG and MEG provides significantly higher spatial resolution for source estimation that either of these modalities alone (Hauk, Stenroos, & Treder, 2019; Henson, Mouchlianitis, & Friston, 2009; Molins, Stufflebeam, Brown, & Hamalainen, 2008). This is essential for studies or applications that require a high degree of anatomical precision, e.g. in connectivity analysis or pattern classification. A few studies have used this combination (mostly with MEG alone) to study saccadic effects in simplified paradigms (Moon et al., 2007; Temereanca et al., 2012), but not during the reading of continuous text.

Here, we simultaneously co-registered eye movements with EEG and MEG while healthy participants read short sentences. Sentence endings could be plausible (“Whe ate the apple”) or implausible (“He ate the shoe”), and participants had to distinguish these categories behaviorally. We could therefore determine at which latency behavioral responses were already affected by contextual information, and which brain responses were likely to correspond to these behavioral effects. We replicated previously reported N400 effects in EEG data during natural reading with our combined EEG and MEG approach, and used their high spatial resolution to determine the spatio-temporal dynamics of the underlying sources. We provide a proof-of-concept that combined EEG and MEG in combination with eye-tracking is feasible and may become a valuable tool to study language and cognition in naturalistic paradigms.

## Methods

### Participants

EEG and MEG data from 18 healthy participants (11 females, average age 28 years, right-handed according to self-reports) were analyzed. Participants had normal or corrected-to-normal vision, and were native speakers of English. They reported no history of reading deficits or neurological/psychiatric disorders. The study was approved by the Cambridge Psychology Research Ethics Committee.

### Stimuli

Material in the plausibility judgement (PJ) task consisted of 248 short sentences in the format [Pronoun] [Verb] [Preposition] [Noun (singular)]. Sentences were derived from 62 quadruples with similar verbs and prepositions but different pronouns and nouns. The noun was thereby altered so that two sentences in a quadruple showed a semantically coherent verb noun relation (plausible sentences) while the remaining two sentences were semantically incoherent in their verb-noun relation (implausible sentences). We present examples in table 1. This design resulted in overall 124 plausible and 124 implausible sentences. In addition, pronouns were also altered within quadruples to increase variability and thereby reduce repetition and prediction effects, while at the same time keeping the stimulus set well-controlled.

**Table 1.** Example Sentence Quadruple

| Pronoun | Verb | Preposition | Noun | Plausibility |
| --- | --- | --- | --- | --- |
| I | frighten | the | calf | Plausible |
| You | frighten | the | class | Plausible |
| They | frighten | the | towel | Implausible |
| We | frighten | the | corn | Implausible |

**Table 2:** Stimulus parameters

| **Table 2:** Stimulus parameters |  |  |  |  |  |
| --- | --- | --- | --- | --- | --- |
|  | Number of<br>letters | Word form<br>frequency<br>p/m | Orthographic<br>neighborhood<br>size | Cloze<br>probability | Plausibility |
| Plausible | 4.8 | 417 | 5.7 | 0.0 | 6.4 |
| Implausible | 5.0 | 383 | 5.7 | 0.0 | 1.9 |

Between quadruples, verbs were not repeated and each noun was only used once in the stimulus set. Classification of sentences into the plausible and implausible categories was achieved on the basis of a plausibility rating of the material, performed with 12 participants that did not participate in the EEG/MEG study. Ratings were obtained on a scale between 1 (not plausible at all) and 7 (highly plausible). Plausible sentences had an average rating of 6.4 (min/max 5.0/7.0), and implausible ones of 1.9 (1.0/3.5).

Sentence final nouns were matched between plausible and implausible conditions for number of letters, bi-gram and tri-gram frequency, orthographic neighbourhood size, imageability and concreteness. Matching was achieved on the basis of the English Lexicon Project database (Balota et al., 2007), the MCWord database (Medler & Binder, 2005) as well as the MRC Psycholinguistic database (Coltheart, 1981). Cloze Probability (Bloom & Fischler, 1980) was close to zero for all sentences, ensuring that final nouns could not be predicted based on preceding context. Cloze probability was zero for all implausible sentences, and different from 0 (equal to 1) for three plausible ones. The cloze probability rating was performed with 14 participants who did not participate in the EEG/MEG study. This control was of particular importance to assure that judgements about the sentences plausibility were only possible after the whole sentence was read.

Stimulus sentences were then split up into four different stimulus lists, each containing one sentence of each quadruple and balanced with respect to the number of plausible and implausible sentences, resulting in 62 sentences per list.

Practice material consisted of stimuli similar, but not identical, to the material of the aforementioned experimental conditions. 20 sentences in the same format as used in the actual experiment where presented to familiarize participants with the plausibility judgement task. These training sentences where balanced for pronouns and plausibility and none of the verbs or nouns used here were later on repeated in the experimental blocks. The practice material also contained 5 sentences that asked the participant to fixate a red or green response field (“Please fixate the red/green triangle”), respectively. These trials allowed the participant to get used to the eye tracking procedure.

### Procedure

MEG, EEG and eye tracking data were acquired in an acoustically and magnetically shielded MEG chamber at the MRC Cognition and Brain Sciences Unit in Cambridge, UK. EEG and MEG signals were recorded with a 306-channel Neuromag Vectorview System (Elektra, AB, Stockholm), containing 204 planar gradiometers and 102 magnetometers, and 70 EEG electrodes mounted on an electrode cap according to the extended 10-20 system (Easycap, reference electrode on nose, ground electrode on cheek). Horizontal and vertical EOG signals were recorded from additional bipolar electrodes. MEG and EEG signals were sampled at 1000 Hz. The position of five Head Position Indicator (HPI) coils, attached to the EEG cap, were digitized with a 3Space Isotrak II System (Fastrak Polhemus Inc., Colchester, VA) in order to determine the head position within the MEG helmet. Three anatomical landmark points (nasion and preauricular points) as well as 50-100 additional randomly distributed points (head shape) were digitized for an accurate co-registration with MRI data.

Eye tracking was conducted using a remote monocular eye-tracker (iView X MEG250, SensoMotoric Instruments, Teltow, Germany) with a sampling frequency of 500 Hz focused on the participant’s dominant eye. Stimulus presentation was controlled using E-Prime software (Version 2, Psychology Software Tools Inc.,Pittsburgh, USA). Stimuli were projected on a screen with dimensions 49 cm by 37 cm which was positioned 130 cm in front of the participant, with a screen resolution of 1024 by 768 pixel, resulting in a visual angle of 0.0208 degree per pixel on screen.

Each participant got written and verbal instructions about the study at the beginning of the experiment. At the beginning of each block the eye-tracker was calibrated using a standard 13 point calibration grind on screen. Calibration was repeated until the validation of this calibration yielded an average measurement error of less than 0.5 visual degree in X and Y for each eye.

During the whole duration of a trial the background was printed in light grey (RGB: 192, 192, 192) and one triangle, printed in red (RGB values: 237, 28, 36), was positioned at the bottom right corner of the screen and one green triangle (RGB values: 181, 230, 29) was presented at the upper right corner of the screen or vice versa. The triangles served as response fields for the experimental tasks and their positions were not switched between trials but between first and second half of the experiment.

For the PJ task, a fixation cross vertically centered on the left hand side of the screen had to be fixated in order to start the presentation of a sentence. Once the fixation cross had been fixated for 1000 ms, the sentence appeared at the right hand side of the fixation cross. The sentence was presented 110 pixels (corresponding to a visual angle of 4.9 degrees) to the right of the fixation cross, and the subsequent words in the sentence were displayed with normal spacing between words. Participants were asked to read the sentence in one go from left to right. Participants were instructed to read at their own pace, but also to indicate the plausibility of the sentence as quickly and accurately as possible by fixating the green or red triangle associated with yes and no responses, respectively. The stimulus sentence and the left fixation cross disappeared when a response was recorded and were replaced by the initial central fixation cross.

### Eye tracking acquisition and pre-processing

Eye-tracking (ET) data were acquired using an MEG-compatible remote SMI tracking device in monocular mode at a sampling frequency of 250 Hz. ET data were processed by an adaptive algorithm (Nystrom & Holmqvist, 2010). Behaviour of subjects was characterized by their reading speed (RS), saccade amplitude (SA), fixation duration (FD), total number of saccades (TNS) and percentage of regressive saccades (PRS). Trials were only included into the analysis if the sentence was correctly classified as plausible or implausible, there were no regressive saccades with amplitudes larger than 1 visual degree, and there was only one fixation on the final target word.

### EEG-MEG pre-processing

Artefacts likely to be produced by sources distant to the sensor array were removed by means of the signal-space separation (SSS) method, which is implemented in the Neuromag “Maxfilter” software (Taulu & Kajola, 2005). The spatio-temporal variant of Maxfilter as well as movement compensation were applied. EEG and MEG data were visually inspected for bad channels, which were marked for interpolation. Data were band-pass-filtered offline between 0.6 and 40 Hz. EEG/MEG recordings were cleaned using independent component analysis (ICA) in MNE-python software. ICA components reflecting eye-movements were determined by correlating their time courses with the eye-tracking signals.

### Source estimation

Source estimates were derived from combined EEG and MEG data using dynamic statistical parametric mapping (dSPM) as implemented in the MNE-Python software package, (Gramfort et al., 2013)) in combination with the FreeSurfer image analysis suite, http://surfer.nmr.mgh.harvard.edu/). The noise covariance matrices for each data set were computed for baseline intervals of 300 ms duration before the onset of sentence presentation. For regularization, the default signal-to-noise ratio was used (SNR=3). EEG/MEG sensor configurations and MRI images were co-registered based on the matching of about 50-100 digitized points on the scalp surface.

High-resolution structural T1-weighted MRI images of each individual subject were used, acquired in a 3 T Siemens Tim Trio scanner at the MRC Cognition and Brain Sciences Unit (UK) with a 3D MPRAGE sequence, field-of-view 256 mm × 240 mm × 160 mm, matrix dimensions 256×240×160, 1 mm isotropic resolution, TR=2250ms, TI=900 ms, TE=2.99 ms, flip angle 9°. The scalp surface of the MRI images was reconstructed by using the automated segmentation algorithms of the FreeSurfer software. By using the traditional method for cortical surface decimation, the original triangulated cortical surface (consisting of several hundred thousand vertices) was down-sampled to a grid with an average distance between vertices of 5 mm, which resulted in approximately 1.000 vertices. A boundary element model (BEM) containing 5120 triangles was created from the inner skull surface for MEG measurements, by using a watershed algorithm. For EEG, the BEM model was created with three layers for scalp, outer skull surface, and inner skull surface. Dipole sources were assumed to be perpendicular to the cortical surface.

Source estimates were computed for each subject and sentence condition. The results for each individual data set were morphed to the average brain across all subjects.

Permutation-based whole-brain cluster-level statistics were applied to the averaged distributions within each time window of interest. A non-parametric cluster-level t-test for spatio-temporal data was used (Maris & Oostenveld, 2007). The permutation test corrected for multiple comparisons across all vertices of whole-brain source estimates within each of the a priori defined time windows.

## Results

Up to the target word, behaviour was similar between plausible and implausible sentences on several relevant variables (Table 3), indicating that our stimulus material was well-matched. Including the target words, subjects read plausible sentences significantly faster (P<0.001) compared to implausible sentences, and had significantly larger TNS (P= 0.0018), PRS (P= 0.0033) and SA (P= 0.0057) for implausible sentences (Table 4). The latter trend was also observed numerically for FD, but the difference did not reach the level of significance (P=0.2953).

**Table 3:** Behavioural results up to the target word.

| **Table 3: Behavioural results up to the target word.** |  |  |  |
| --- | --- | --- | --- |
| **Measure** | **Plausible** | **Implausible** | **P** |
| Reading speed<br>(words/s) | 5.43 | 5.72 | n. s. |
| Total number of<br>saccades | 350.5 | 345.5 | n. s. |
| Percentage of<br>regressive saccades<br>(%) | 1.82 | 1.91 | n. s. |
| Saccade amplitude<br>(vis. deg.) | 2.99 | 3.04 | n. s. |
| Fixation duration (s) | 0.126 | 0.130 | n. s. |

**Table 4:** Behavioural results for target words.

| **Table 4:** Behavioural results for target words. |  |  |  |
| --- | --- | --- | --- |
| **Measure** | **Plausible** | **Implausible** | **P** |
| Reading speed<br>(words/s) | 3.93 | 3.65 | <0.001 |
| Total number of<br>saccades | 436 | 463 | <0.01 |
| Percentage of<br>regressive saccades<br>(%) | 3.65 | 5.71 | <0.01 |
| Saccade amplitude<br>(vis. deg.) | 2.92 | 2.94 | <0.01 |
| Fixation duration (s) | 0.150 | 0.152 | n. s. |

ICA was successful at reducing the effect of eye-movements on both EEG and MEG signals (Figure 1). Visual inspection of fixation-related EEG/MEG data revealed event-related potential components around 90 and 250 ms in occipital and occipito-temporal sensors as well as another component with a broader time range (270-460 ms) in centro-parietal sensors (Figure 2). The latter component showed more negative-going deflection for the implausible words. In order to test for the presence of an N400-type response in our EEG data, we ran consecutive paired t-tests on ERP signals from electrode Cpz. The comparison was significant at an uncorrected level between about 200 and 400 ms, and at false-discovery-rate-corrected level around 270 ms and 330 ms, with signals for implausible sentences being more negative-going (Figure 2).

**Figure 1:**
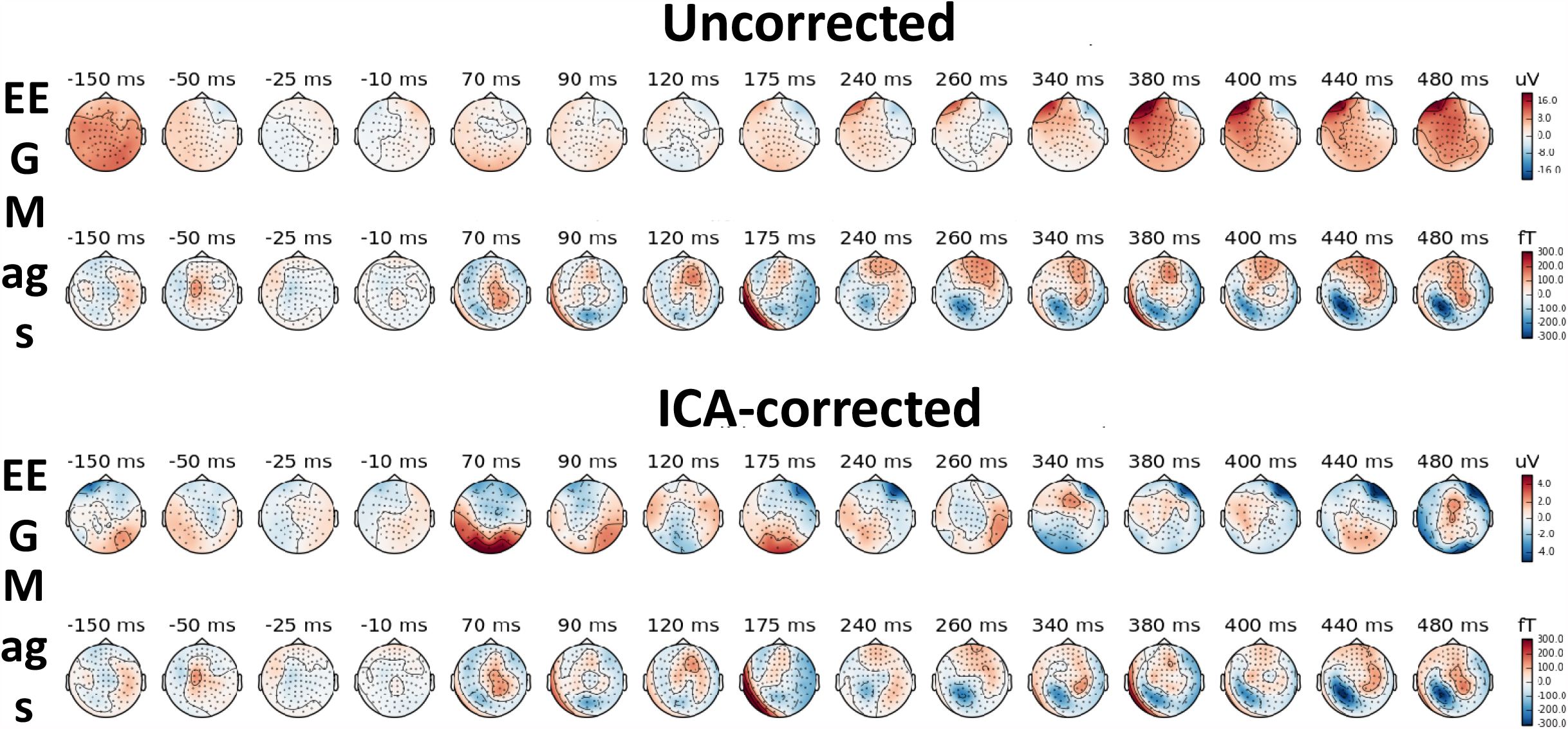
Signal-space topographies for fixation-related potentials (EEG) and fields (magnetometers) for the sentence-final word before (top) and after (bottom) ICA artefact correction. Note the difference in scale between uncorrected and ICA-corrected topographies for EEG.

**Figure 2:**
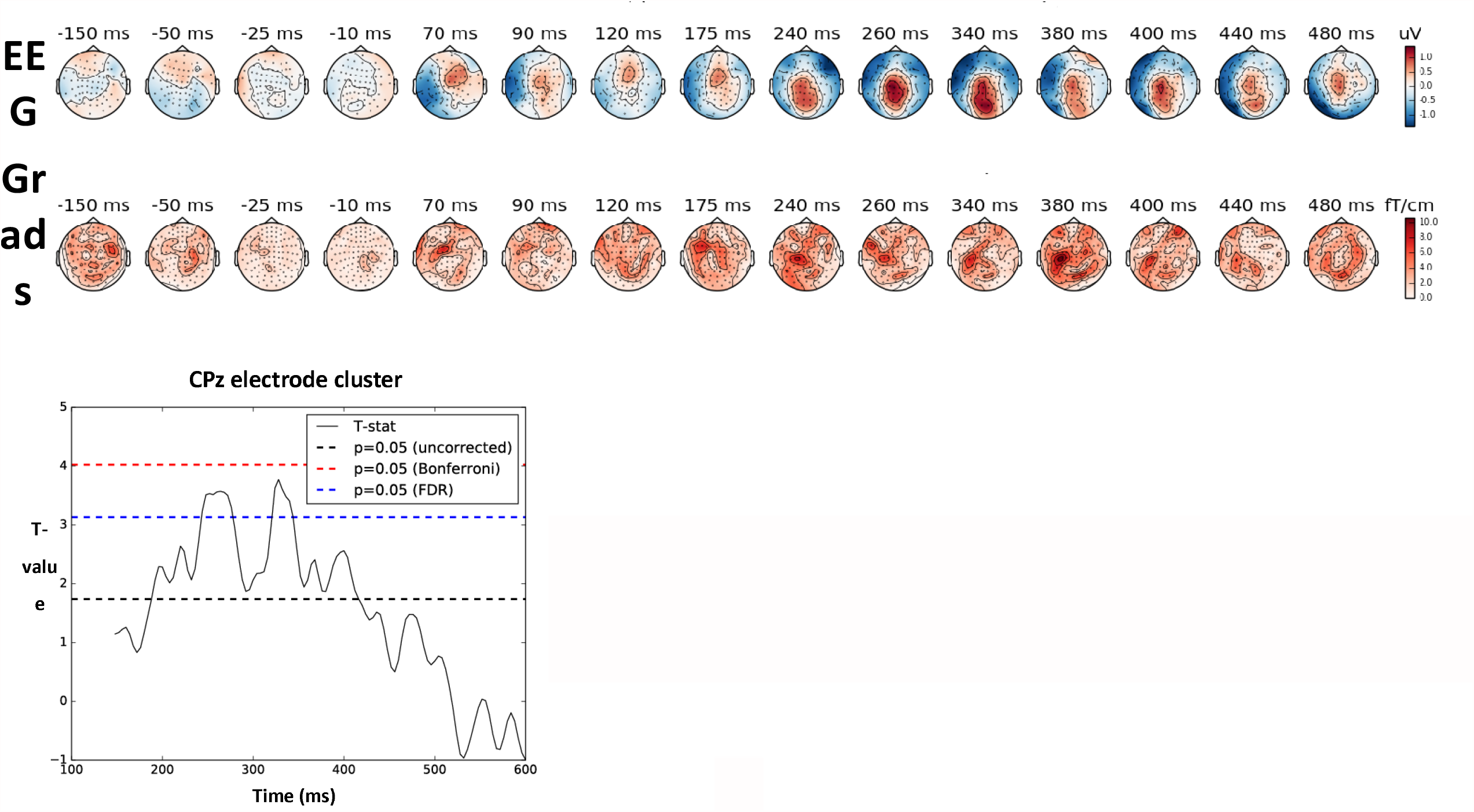
Topographies (top) and statistics (left) for the difference Plausible minus Implausible sentence-final words. The EEG topography after 240 ms resembles the expected N400 (more centro-parietal negative-going potential for Implausible sentences).

L2-minimum-norm estimates for all target words (averaged across plausible and implausible sentences) produced activity in occipital and left-temporal brain regions (Figure 3). Cluster-based permutation testing resulted in a significant cluster (P=0.005) comprising medial temporal, inferior frontal and supplementary motor brain regions in the left hemisphere, with stronger activations for plausible compared to implausible words (Figure 4).

**Figure 3:**
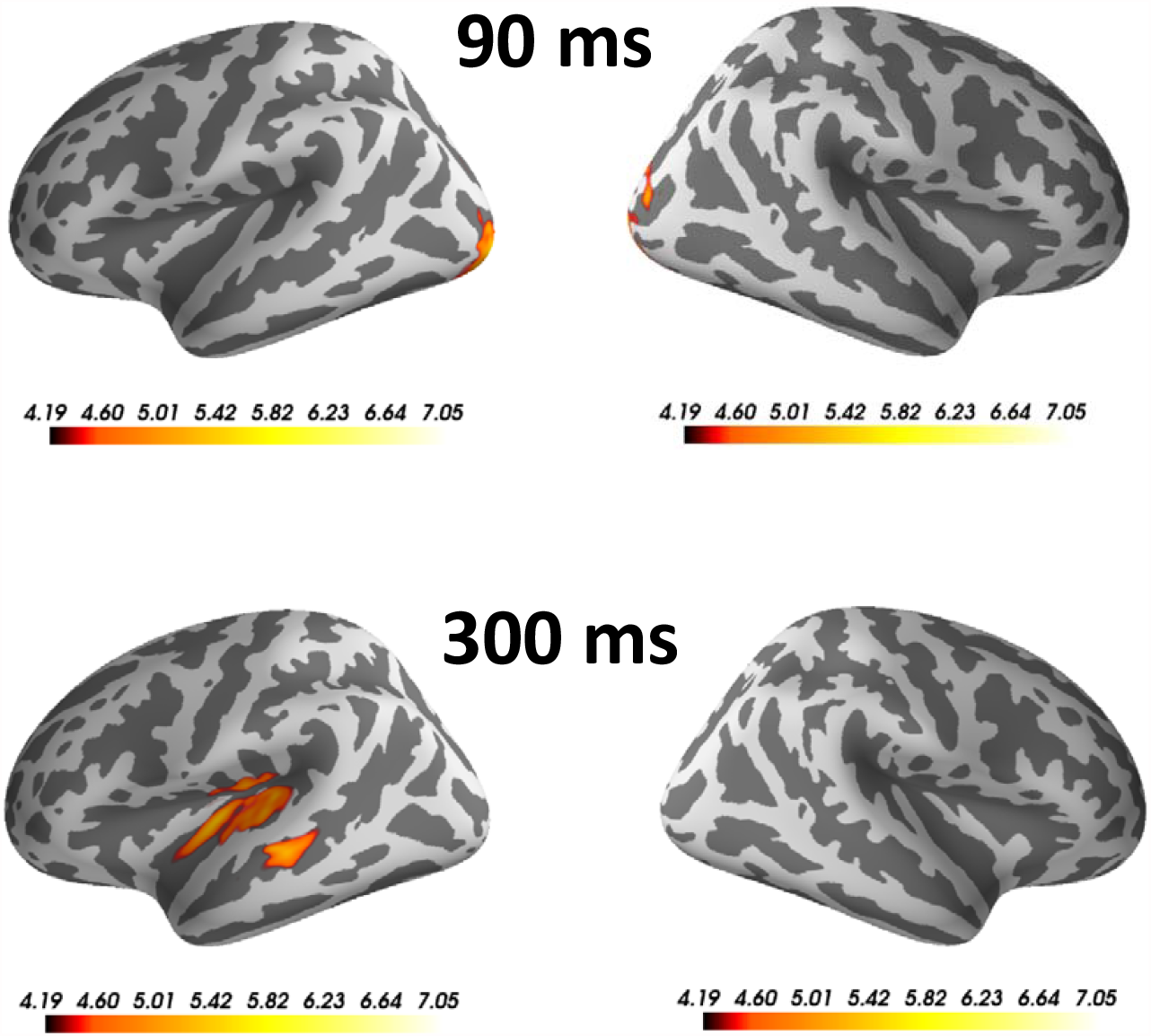
L2-minimum-norm source distributions from combined EEG and MEG data for the average across all target words, at different latencies.

**Figure 4:**
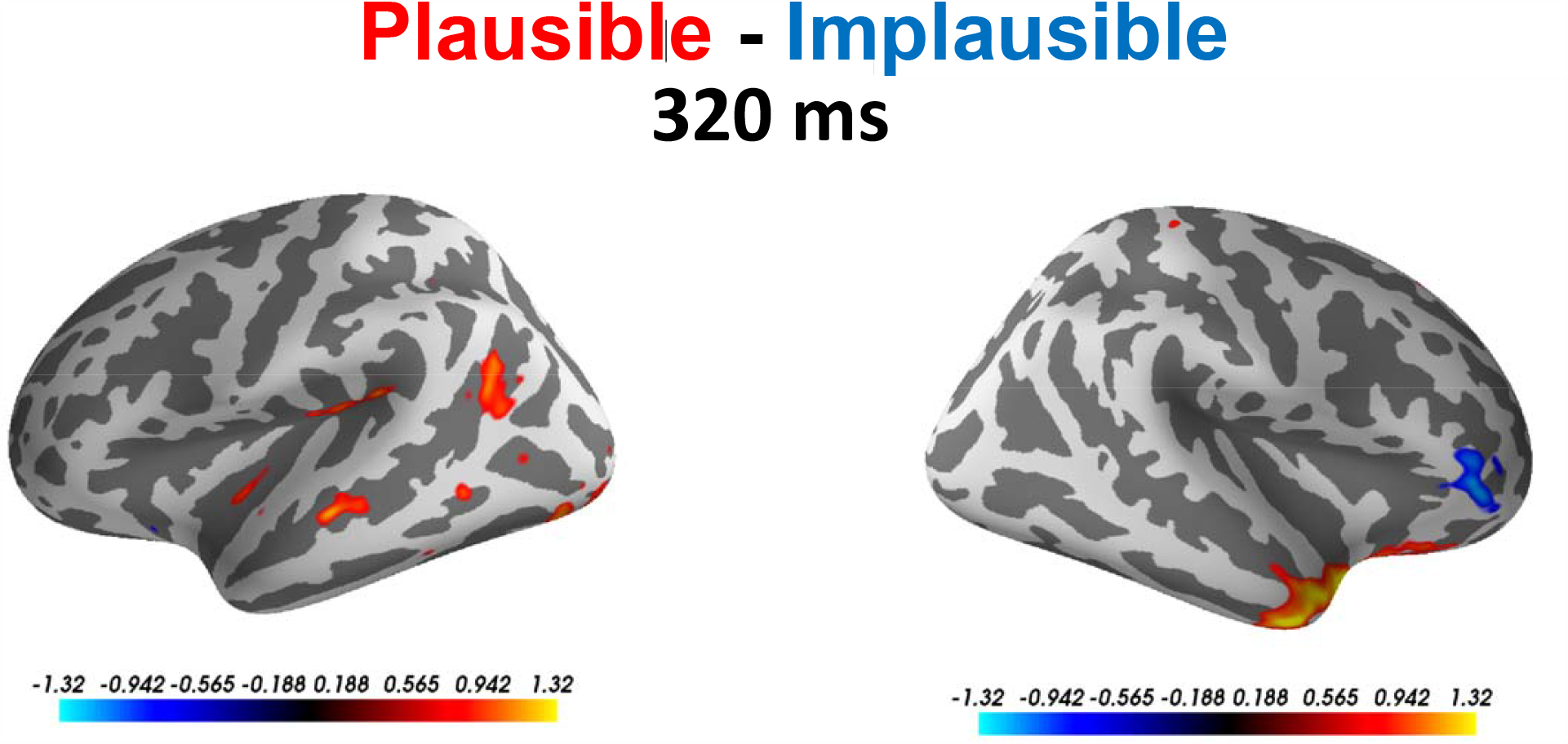
Difference between source distributions for Plausible and Implausible target words at 320 ms.

## Discussion

We provided a proof-of-concept that the co-registration of eye-tracking and combined EEG/MEG is feasible and allows the estimation of brain activity underlying natural reading processes. We replicated the well-known N400 effect in a simple but naturalistic sentence reading task, and obtained plausible source estimation results from combined EEG/MEG data. This methodology may in the future contribute to the deeper understanding of the neural mechanisms of natural reading, and higher cognitive function more generally.

The co-registration of eye-tracking and EEG has been used in a number of previous studies (e.g. Dimigen et al., 2011; Himmelstoss et al., 2019; Hutzler et al., 2007; Kornrumpf et al., 2016), e.g. to study specific effects of para-foveal preview (Degno et al., 2019; Dimigen, Kliegl, & Sommer, 2012) and text format (Weiss et al., 2016). Eye-tracking has also been used to investigate natural reading processes in fMRI (Desai et al., 2016; Schuster et al., 2016). The co-registration of eye-tracking with combined EEG and MEG will enable us to study brain processes at the behavioural and neural level with both high temporal and reasonable spatial resolution. The combination of spatial and temporal resolution is required to unravel the brain networks supporting natural language processing (Hauk, 2016). New developments in non-cryogenic on-scalp MEG sensors may further improve the sensitivity and spatial resolution of EEG/MEG methodology (Boto et al., 2018; Iivanainen, Stenroos, & Parkkonen, 2017).

In our present study, we made a first step in this direction. We compared four-word sentences with plausible and implausible final words. Of course, in the future we would like to use more naturalistic stimuli, i.e. longer sentences, paragraphs or stories. This would also allow investigating regressions during reading of natural text (e.g. Booth & Weger, 2013), a behaviour that cannot be studied using conventional word-by-word or single-sentence paradigms. We used ICA to remove eye-movement-related artefacts from our EEG/MEG data. This potentially removes activity from brain sources whose time courses are correlated with those of eye movements. In the future, artefact-correction algorithms based on signal-space projections using explicitly measured artefact topographies (Dimigen et al., 2011; Ille, Berg, & Scherg, 2002) may increase sensitivity to some psycholinguistic variables.

Our methodology opens new avenues to study the brain networks supporting language and other domains of cognition in more detail. Reichle et al. (2011) sketched out the possible network underlying eye movement control during reading. However, their proposal does not contain several brain regions in temporal and frontal lobes known to be involved in semantic information retrieval and semantic control (Binder, Desai, Graves, & Conant, 2009; Pulvermuller, 2013; Ralph, Jefferies, Patterson, & Rogers, 2017; Taylor, Rastle, & Davis, 2013). At a typical reading speed of four words per second, a 30-minute experiment could potentially deliver behavioural and brain data for 7200 words. These data are sensitive to a large range of perceptual, psycholinguistic and cognitive variables. The richness of datasets based on naturalistic experimental paradigms, such as listening to stories or watching movies, has already been exploited in recent fMRI studies (e.g. Baldassano, Hasson, & Norman, 2018; Huth, de Heer, Griffiths, Theunissen, & Gallant, 2016; Nishimoto et al., 2011; Regev et al., 2018). Methods for analysing large numbers of predictor variables and overlapping brain responses in EEG/MEG data have recently become available (Crosse, Di Liberto, Bednar, & Lalor, 2016; Ehinger & Dimigen, 2018). In the future, this will be useful to characterise the link between brain and behaviour, inter-individual differences in language and cognitive processes, and may lead to diagnostic and prognostic tools for brain diseases.

## Acknowledgements

This work was supported by intramural funding from the Medical Research Council UK (MC_UU_00005/18) and by the MRC UK MEG Partnership Grant (MR/K005464/1). The authors would like to thank Caroline Coutout for her help during the experiment set-up and data acquisition.

## Notes

### Competing Interest Statement

The authors have declared no competing interest.

